# Poor foam stability in gluten-free beers is associated with distinct protein composition compared with barley-malt beers

**DOI:** 10.64898/2026.09.04.749538

**Authors:** Zesun Lyu, Jacob Humphries, Farhad Masoomi-Aladizgeh, Thomas H. Roberts

**Affiliations:** School of Life and Environmental Sciences, University of Sydney, Camperdown, NSW 2006, Australia; Sydney Institute of Agriculture, University of Sydney, Eveleigh, NSW 2015, Australia

**Keywords:** foam stability, gluten-free beer, proteomics, serpin, lipid transfer protein

## Abstract

**Why was the work done?:** Gluten-free (GF) beers typically exhibit poorer foam stability than conventional barley malt-based beers, yet the molecular basis underlying this difference remains poorly understood. This study aimed to investigate protein-level factors associated with foam stability in GF and barley beers.

**How was the work done?:** Three commercially produced GF beers, comprising two lagers brewed from rice or sorghum, and one pale ale brewed from a millet-buckwheat-rice blend, were compared with two commercially produced barley-malt beers, comprising a lager and a pale ale. Soluble protein was isolated from freeze-dried beer powder using TCA-acetone. Protein profiles were examined using SDS-PAGE, and proteomes were characterised and compared using label-free quantitative LC-MS/MS.

**What are the main findings?:** GF beers showed lower soluble protein concentrations and significantly lower foam stability than the barley beers. SDS-PAGE revealed fewer and weaker protein bands in GF beers compared with barley beers. Proteomic analysis showed that barley beers were enriched in known foam-active proteins, particularly lipid transfer protein 1 (LTP1) and Protein Z (serpin family members), whereas these proteins were not detected or were detected only at very low levels in GF beers.

**Why is the work important?:** These findings indicate that reduced soluble protein abundance and the absence or low abundance of barley-associated foam-active proteins are key proteomic features distinguishing GF beers from conventional barley-based beers. This work improves current understanding of foam quality in GF beer and provides a foundation for future studies aimed at improving foam stability.

## Introduction

Beer foam stability is an important quality parameter in brewing, contributing to the visual appearance, mouthfeel, and overall perception of beer quality (Habschied et al. 2022). In brewing, a fine, uniform, and persistent foam is widely regarded as a hallmark of high-quality beer, and its consistency is therefore a key concern for brewers during recipe formulation and process control (Lewis and Young 2001). The formation and retention of beer foam depend on the stability of the gas–liquid interface, which is strongly influenced by foam-active components derived from cereal raw materials (Lu et al. 2020). Although factors such as carbonation level, hop bitter acids, and polysaccharides may contribute to foam behaviour, proteins with surface-active properties play a central role in stabilising the foam film and maintaining its elasticity (Evans and Hejgaard 1999; Caballero et al. 2012; Viejo et al. 2019; Reid et al. 2024).

In conventional barley malt beers, lipid transfer protein 1 (LTP1), a low-molecular-weight protein, and Protein Z, which represents members of the serpin family (Hejgaard et al. 1985), are widely recognised as the principal foam-positive proteins (Evans et al. 2003). LTP1 readily adsorbs to the gas–liquid interface during pouring and early foam development, facilitating foam formation, whereas Protein Z contributes to foam retention by enhancing the structural integrity and elasticity of the foam film (Evans et al. 2003, Perrocheau et al. 2006). Together, these proteins form a complementary system that governs both the generation and stability of beer foam in traditional lagers and ales.

In contrast to conventional beers, gluten-free (GF) beers, which are brewed from alternative cereals such as rice, sorghum, buckwheat, or millet, typically exhibit inferior foam quality, characterised by poor foam formation and rapid foam collapse (Cela et al. 2023). As the demand for GF beverages continues to grow along with the market growth of GF products generally (Maior et al. 2024), these deficiencies represent a persistent challenge for brewers, as foam quality is a key determinant of product appearance and consumer acceptance (Habschied et al. 2022).

One potential explanation for the disparity in foam behaviour between barley-based and GF beers lies in fundamental differences in grain protein composition. In particular, the presence, abundance, and functionality of foam-relevant proteins in GF beers remain poorly characterised compared with those in barley-based beers. Previous studies on GF beer production have often focused on process optimisation and broader quality attributes rather than directly examining how protein composition relates to foam performance. Although the roles of LTP1 and Protein Z have been extensively characterised in barley beer systems, it remains unclear whether functionally analogous proteins are present in GF cereals, or how their abundance and behaviour in finished beer relate to foam stability.

To address this knowledge gap, the present study combines standard foam stability measurements with liquid chromatography–tandem mass spectrometry (LC–MS/MS)-based proteomic analysis to compare the foam properties and protein profiles of gluten-containing and GF beers. By examining differences in protein composition across barley-based and alternative-grain beers, this work aims to provide insight into the molecular basis underlying reduced foam stability in GF beer.

## Materials and methods

### Materials and sample preparation

Five commercially produced beers, comprising two gluten-containing controls and three GF samples, were purchased from local retail outlets in Sydney, Australia, on two separate occasions, 1 month apart. The gluten-containing beers consisted of a barley-malt lager and a barley-malt pale ale, both selected as conventional barley malt-based beers based on product labelling. The GF samples consisted of a rice-based lager, a sorghum-based lager, and a mixed-grain pale ale made from millet, buckwheat, and rice. Publicly available product information indicated the use of malted grains in the mixed-grain GF beer, although it was not possible to confirm whether all grain components were malted. The malting status of the grains used in the rice- and sorghum-based beers could not be verified because manufacturers declined to disclose recipe details. Immediately after opening, beer samples were transferred into 50-mL Falcon tubes (pre-cooled in a −80°C freezer), flash-frozen at −80°C for 30 min and subsequently freeze-dried using an ALPHA 1-4 LSCbasic freeze dryer (Gefriertrocknungsanlagen GmbH, Germany). Freeze-drying was conducted with a pre-freezing temperature of −60°C, a chamber pressure of ≤0.1 mbar, and a total drying duration of 48 h. The resulting lyophilised powders were sealed and stored at −80°C until further analysis.

### Protein extraction and quantification

Protein extraction was performed using a modified trichloroacetic acid (TCA)-acetone precipitation approach adapted from Rinaldi (2023). Preliminary extraction trials compared conventional TCA-acetone precipitation, in which TCA was dissolved in cold acetone, with a modified aqueous TCA precipitation approach, in which TCA was first dissolved in ddH₂O before acetone washing. The modified approach was informed by Ngo et al. (2014), who reported that TCA-mediated protein precipitation involves the formation of a partially folded, sticky aggregation-like “A-state” and that lower final concentrations of aqueous TCA can improve protein precipitation from low-protein samples. During optimisation, the modified aqueous TCA protocol produced a higher level of protein extraction from GF beer samples than the conventional TCA-acetone method, as indicated by bicinchoninic acid (BCA) assay. This protocol was therefore selected for the final extraction workflow and applied to all beer samples to minimise methodological variation.

Approximately 100 mg of freeze-dried beer powder from each sample was suspended in 100 µL of deionised water and transferred to a 2-mL microcentrifuge tube. Samples were vortex-mixed thoroughly and centrifuged at 15,900 × g for 20 min at 4°C. The supernatant, containing the soluble protein fraction, was retained and mixed with 400 µL of ice-cold 20% (w/v) TCA prepared in ddH₂O to precipitate proteins. The mixture was incubated at −30°C for 10 h and centrifuged as above. The resulting protein pellet was washed three times with pre-chilled 80% acetone. After each wash, samples were vortexed briefly and centrifuged at the same g force for 5 min. Pellets were air-dried at room temperature until residual acetone was removed and subsequently dissolved in 8 M urea in 100 mM Tris-HCl (pH 8.0).

For the GF samples, duplicate extractions were prepared from the beer and then combined during the urea reconstitution step. The combined pellets were dissolved in the urea buffer, vortexed thoroughly, and subjected to sonication using a sonication bath for up to 1 min. The sample was centrifuged at 10,000 × g for 5 min, and the supernatant collected as the protein extract.

Protein concentration was determined using a BCA assay kit (Thermo Fisher Scientific, USA) with bovine serum albumin (BSA) as the standard, following the manufacturer’s instructions. Absorbance was measured at 562 nm using a CLARIOStar microplate reader (BMG Labtech, Germany). A single protein extract from each beer sample was assayed in triplicate (three wells in the 96-well plate), and the mean value was used for subsequent analyses.

### SDS-PAGE of soluble beer proteins

Protein profiles were analysed by SDS-PAGE under reducing conditions. Protein extracts were mixed with 2× Laemmli sample buffer and dithiothreitol (DTT), giving a final DTT concentration of 32 mM, followed by heating at 95°C for 10 min. Samples were separated on 4–20% Mini-PROTEAN TGX precast gels (Bio-Rad, USA) using 1× Tris/Glycine/SDS running buffer at 120 V. Loading volumes were 7 μL for gluten-containing samples and 15 μL for GF samples to account for the lower protein concentration of the GF beers. A Precision Plus Protein™ Kaleidoscope Standards ladder (Bio-Rad, USA) was used as the molecular weight marker. Gels were stained with Coomassie Brilliant Blue R-250 (0.1% w/v in 40% methanol and 10% acetic acid) and destained until a clear background was achieved. Gel images were acquired using a ChemiDoc MP Imaging System (Bio-Rad, USA).

### In-solution digestion

In-solution trypsin digestion was performed using sequencing-grade modified trypsin (Promega, USA) as described previously by Masoomi-Aladizgeh et al. (2021). Briefly, protein extracts were diluted in 100 mM Tris–HCl (pH 8.0). DTT and iodoacetamide were then added to final concentrations of 10 mM and 20 mM, respectively, with reduction performed at room temperature for 30 min and alkylation at room temperature in the dark for 30 min. Trypsin was added to each sample at 1 µg total proteinase per sample, corresponding to 2 µL of a 0.5 µg/µL stock solution, and samples were incubated at 37°C overnight. Digestion was terminated by adding formic acid to a final concentration of 1% (v/v). Peptide digests were desalted using Sep-Pak C18 cartridges (Waters, USA) according to the manufacturer’s instructions and then vacuum-dried and reconstituted in 0.1% formic acid.

### Sequential window acquisition of all theoretical fragment ion spectra (SWATH-MS)

Peptides were analysed on a Shimadzu LC-40D X3 coupled to a ZenoTOF 7600 mass spectrometer (SCIEX, USA). A 10-µL aliquot of the reconstituted peptide digest was injected at 0.8 mL min⁻¹ using an 8-min gradient, with the autosampler held at 8°C and column oven at 40°C. Mobile phase A was water with 0.1% formic acid, while mobile phase B was acetonitrile with 0.1% formic acid. Data were acquired in positive electrospray ionisation (ESI) mode using Zeno SWATH data-independent acquisition (DIA). A TOF-MS scan (*m/z* 400–900, 0.01 s) was followed by sequential MS/MS acquisition over *m/z* 100–1800 using collision-induced dissociation (CID) with a 0.005 s accumulation time per window. Blank injections were included between samples, and SWATH-MS runs were acquired in block-randomised injection order.

### Proteomics analysis

Data from the DIA mode were processed using MaxQuant (version 2.7.4.0) with default parameters against the UniProtKB database for *Hordeum vulgare* (130,189 sequences), *Fagopyrum esculentum* (745 sequences), *Pennisetum glaucum* (118,277 sequences), *Oryza sativa* (625,414 sequences), and *Sorghum bicolor* (96,443 sequences), released in September 2025. Trypsin/P was specified as the protease with up to two missed cleavages allowed. Carbamidomethylation of cysteine was set as a fixed modification, oxidation of methionine as a variable modification, and peptide and protein identifications were filtered at a false discovery rate (FDR) of 1%.

### Foam stability assay

Foam stability was evaluated according to the American Society of Brewing Chemists (ASBC) Beer– 22 Foam Collapse Rate method (2018), using a narrow outlet 500-mL separatory funnel instead of a 1,000-mL ASBC foam funnel. All glassware was rinsed with deionised water and dried prior to use. Beer samples were equilibrated to 22 ± 1°C and gently poured into the funnel to generate a continuous foam column approximately 400 mL in volume. During drainage, analytical-grade isopropanol (≥99.7%) was applied dropwise at the funnel outlet to collapse foam emerging with the drained liquid, as specified in ASBC Beer–22.

At the end of the collapse period time, t (seconds), the drained liquid volume, b (mL), was recorded. The residual foam remaining in the funnel at time t was allowed to fully collapse, and the additional liquid volume obtained, c (mL), was measured. Foam stability was expressed as the Sigma value (Σ_500_), calculated as:

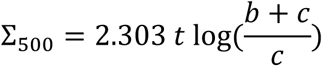

Each sample was analysed in triplicate using freshly opened beer samples, and the funnel was rinsed and dried between replicates.

### Statistical analysis

Statistical analyses were performed in R version 4.5.1. Soluble protein concentrations were compared among beer samples using one-way analysis of variance (ANOVA), followed by Tukey’s post-hoc test for pairwise comparisons. Foam stability indices (Σ_500_) were compared between barley-based and GF beers using Welch’s two-sample t-test. Differences were considered statistically significant at p < 0.05. Data visualisation was performed using ggplot2.

## Results

### Gluten-free beers exhibit markedly lower soluble protein content than barley-based beers

Following optimisation of the extraction procedure, soluble protein concentrations were quantified by BCA assay and compared among beer samples using one-way ANOVA followed by Tukey’s post-hoc test (Figure 1). Both barley-based beers had higher soluble protein concentrations than each of the three GF beers (*p* < 0.001 for all Tukey-adjusted post-hoc comparisons). The barley lager had the highest protein concentration, with 67.4 μg·mL^−1^, and the barley pale ale had 40.5 μg·mL^−1^. In contrast, the GF beers displayed uniformly low protein concentrations, with values ranging from 1.2 to 3.3 μg·mL^−1^, more than an order of magnitude lower than those observed in the barley-based beers. No statistical differences in soluble protein concentration were detected among the three GF beers.

**Figure 1.**
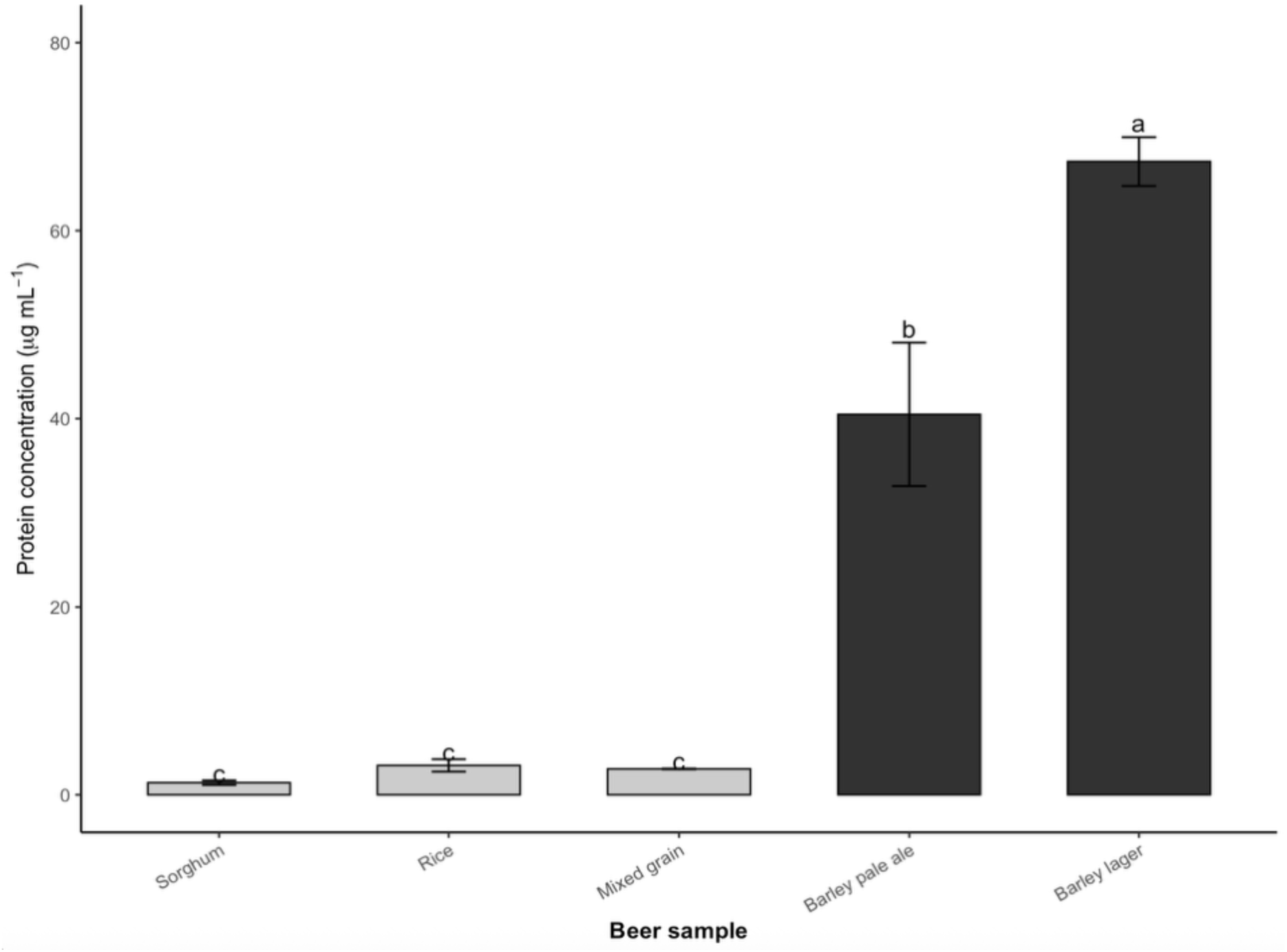

### Protein banding profiles differ between barley-based and gluten-free beers

SDS-PAGE analysis revealed distinct protein banding patterns among the beer samples (Figure 2). The barley-based beers exhibited multiple prominent bands predominantly within the 10–50 kDa range, whereas the GF beers showed fewer and fainter bands on the gel. Overall, these gel patterns were consistent with the markedly higher soluble protein concentrations in the barley-based beers measured by BCA assay.

**Figure 2.**
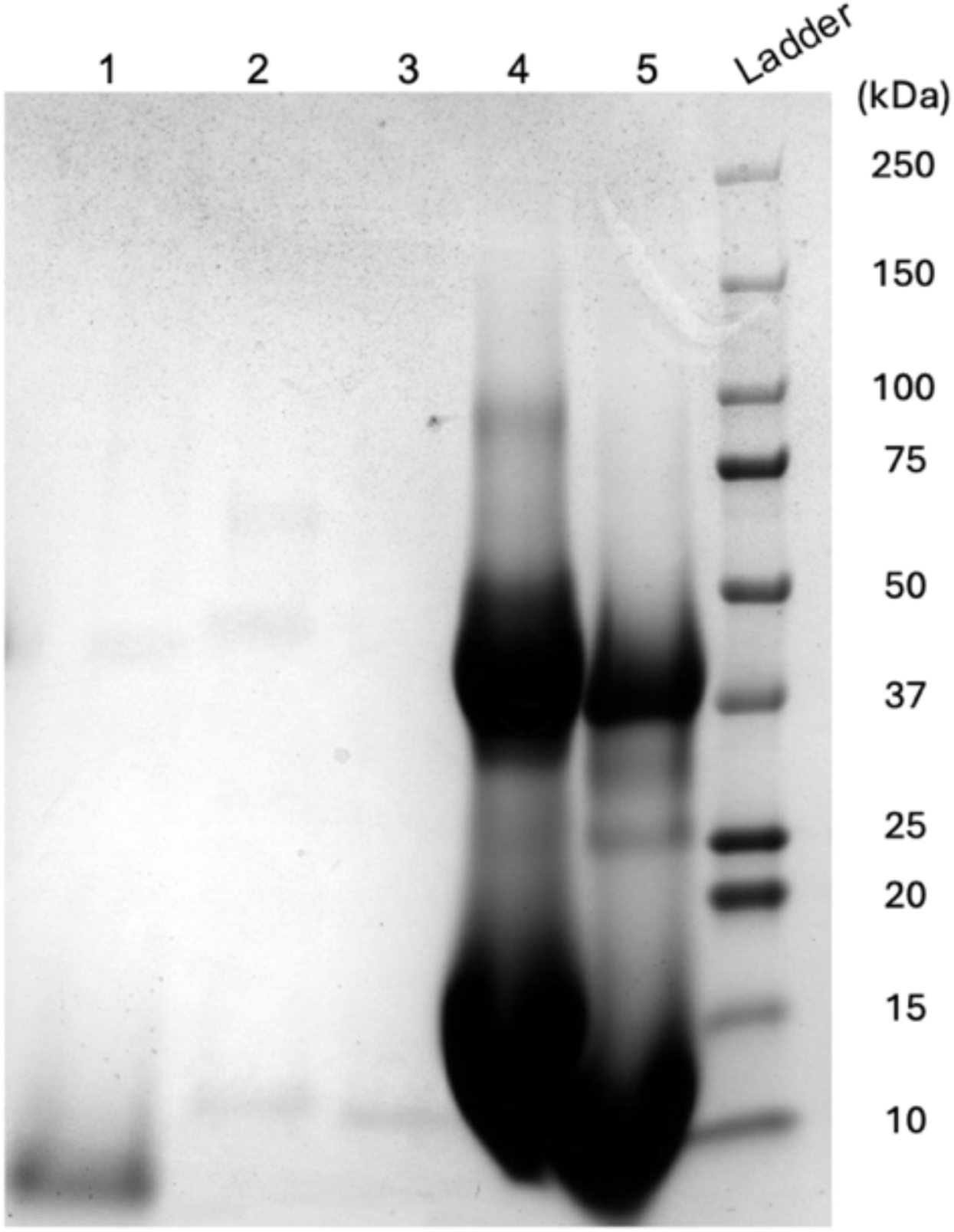

### Gluten-free beers have lower foam stability

The mixed-grain GF beer was excluded from the Σ_500_ analysis because its foam stability was so poor that its foam collapsed too quickly to be measured reliably. The mean Σ_500_ was 110 for conventional beers and 84 for GF beers with a mean difference between groups of 26 (95% CI: 16, 37), indicating poorer foam retention among GF beers (Figure 3).

**Figure 3.**
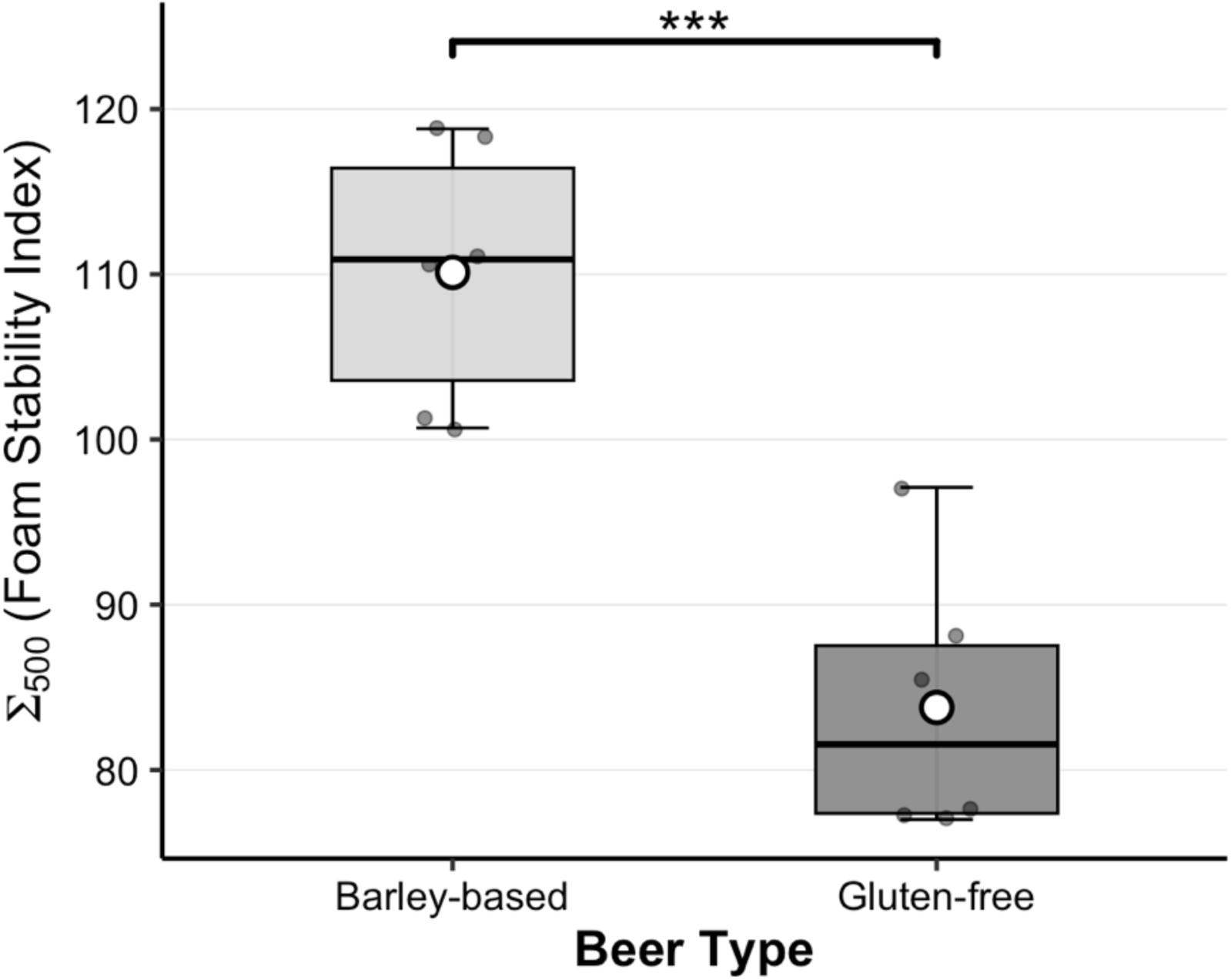

### LC–MS/MS reveals distinct proteomic profiles between barley-based and gluten-free beers

LC–MS/MS analysis revealed marked differences in protein composition between the barley-based and GF beer samples. Proteins were identified using MS/MS database searching in MaxQuant with default parameters. Protein abundances were compared based on signal intensity, and the dominant proteins were grouped according to brewing-relevant functional roles to assess their relative contribution across beer types.

For the barley-based beers, the identified proteins were dominated by storage proteins and protease inhibitors typically associated with barley malt, including hordeins, lipid transfer protein 1 (LTP1), and Protein Z isoforms (Z4 and Z7). In contrast, GF beers brewed from rice, sorghum, and mixed grains lacked the strongly dominating proteins observed in the barley-based beers, and instead displayed proteomes characterised mainly by metabolic enzymes and non-specific LTPs.

To illustrate the compositional differences between the beers, proteins were grouped according to their beer quality-relevant functional roles, and the cumulative signal intensity within each category was used to estimate their relative contribution to the dominant proteome (Figure 4). This intensity representation highlights the strong dominance of foam-related proteins in barley beers, with GF beers displaying a more heterogeneous functional distribution. At the same time, the overall MS signal intensities of the 10 most abundant proteins in GF beers were markedly lower than that observed in the barley-based samples, consistent with the lower soluble protein levels measured by BCA assay. Accordingly, lower signal intensity in the GF beers likely reflects both lower soluble protein concentration and differences in the relative distribution of detected proteins. Detailed identities and intensities of the 10 most abundant proteins detected in each sample are provided in Supplementary Tables S1–S5.

**Figure 4.**
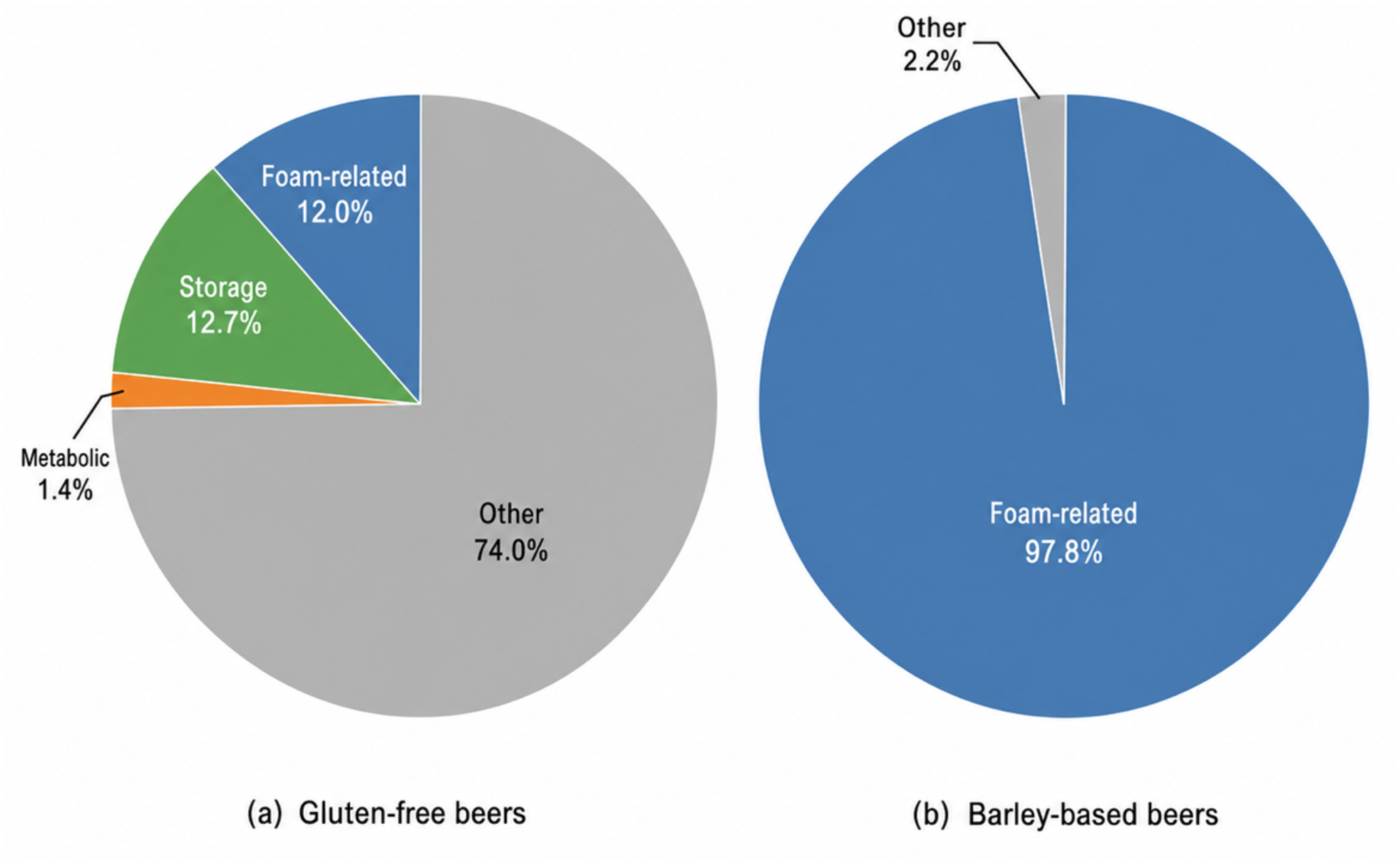

### Foam-active protein families are enriched in barley-based beers

To specifically examine proteins associated with beer foam formation and stability, the relative abundance of representative foam-active protein families was visualised using a comparative heatmap based on log_2_-transformed intensities (Figure 5). The proteins analysed included lipid transfer proteins (LTPs), Protein Z isoforms (Z4 and Z7), and hordeins. Barley-based beers (lager and pale ale) exhibited consistently higher relative abundances of these foam-active protein families compared with GF beers. In particular, LTP1 and Protein Z isoforms were strongly enriched in both barley-based samples but were absent or detected only at very low levels in the GF beers.

**Figure 5.**
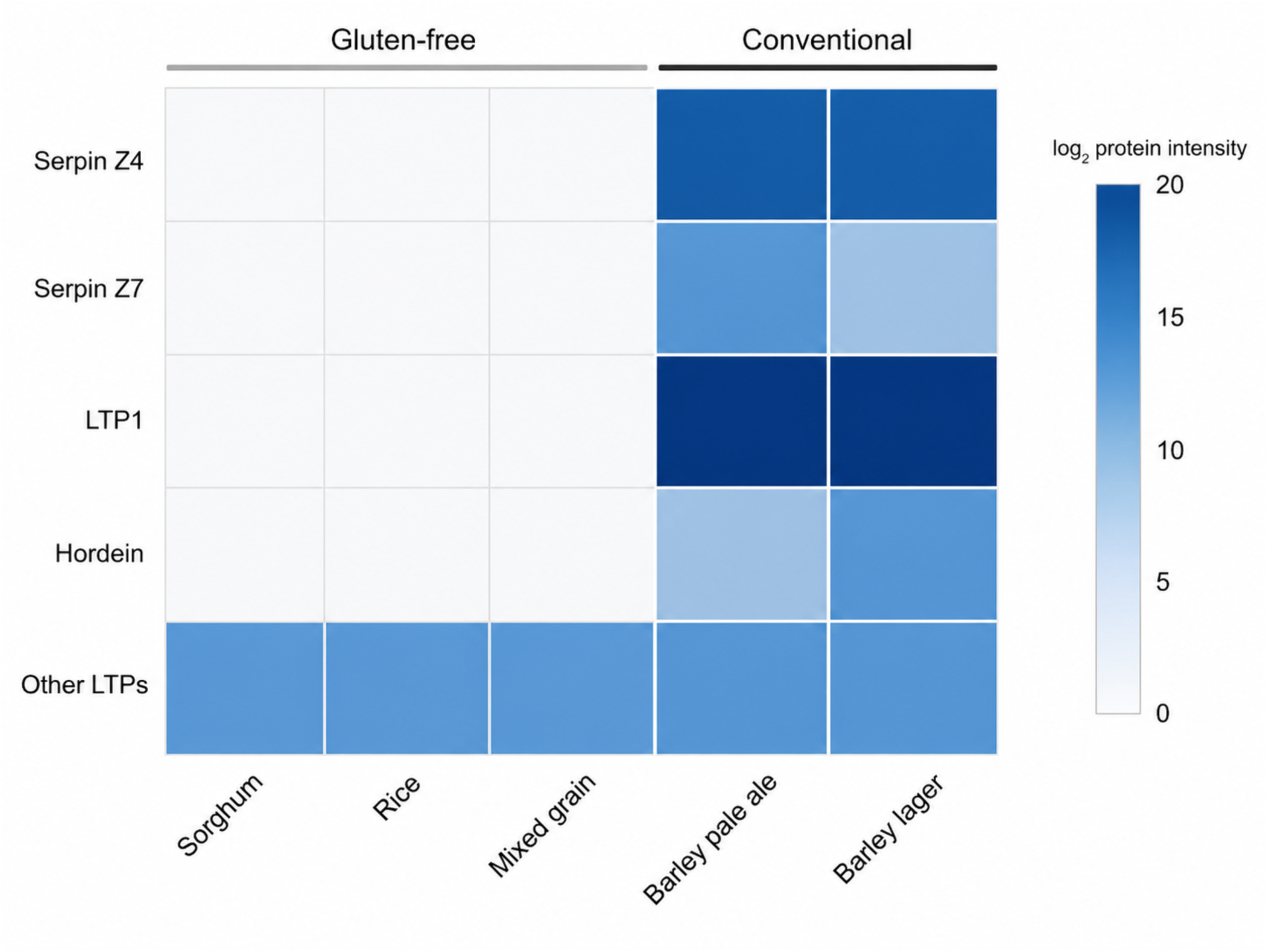

Hordein-derived proteins were exclusively detected in the barley-based beers and were completely absent from all the GF samples, as expected based on raw material composition. In contrast, GF beers showed only low-level signals corresponding to non-specific lipid transfer proteins, with no detectable enrichment of protein families previously associated with foam stability in conventional beer systems.

## Discussion

### Protein composition underpins foam stability differences between barley-based and gluten-free beers

This study demonstrates that differences in foam stability between barley-based and GF beers are closely associated with disparities in soluble protein abundance and composition. Across all samples, barley-based beers consistently exhibited higher soluble protein levels and higher foam stability compared with GF beers brewed from rice, sorghum, and mixed grains. This association reinforces the established view that soluble, surface-active proteins play a central role in the formation and stabilisation of beer foam by constructing a cohesive and elastic interfacial film at the gas–liquid interface (Evans and Hejgaard 1999).

Beer foam stability depends on a dynamic balance between foam-promoting, foam-modifying, and foam-negative components. In conventional barley-based systems, this balance is maintained through the presence of multiple foam-active proteins that contribute both interfacial activity and mechanical reinforcement (Niu et al. 2018; Bamforth 2023). In contrast, the proteomic profiles of GF beers revealed a substantial shift in this balance, characterised by a very low abundance of foam-promoting proteins and a simplified soluble protein repertoire. Such compositional changes are expected to compromise the continuity and elasticity of the interfacial film, predisposing GF beers to rapid foam collapse.

Proteomic analysis further indicated that the soluble protein fraction of GF beers has only minor contributions from proteins previously associated with foam stability in conventional beers. This contrasts sharply with barley-based beers, in which storage proteins, LTPs, and serpin family members collectively constitute a substantial proportion of the soluble proteome. The reduced diversity and abundance of surface-active proteins in GF beers may also limit their overall foam-forming and foam-stabilising capacity, contributing to the lower foam stability observed in these beers. These findings suggest that impaired foam stability in GF beers may result from a broader deficiency in the protein sets required for foam formation and retention.

Taken together, these findings are best interpreted as evidence for a broad alteration in the foam-associated proteome rather than the effect of a single missing protein. Alongside the lower total soluble protein concentrations, the absence or marked reduction in the abundance of key foam-active proteins may compromise the protein contributions required for foam formation and retention, providing a plausible molecular explanation for the consistently lower foam stability observed in GF beers.

### Complementary roles of LTP1 and Protein Z in beer foam formation and stabilisation

In conventional barley-based beers, LTP1 and Protein Z are widely recognised as two major contributors to foam stability. Rather than contributing through the same mechanism, these proteins appear to play complementary roles at different stages of foam development, collectively supporting the foam density and head retention of the foam structure.

### Interfacial activity of LTP1 during early foam formation

LTP1 is a small, highly amphiphilic protein that rapidly migrates to the gas–liquid interface during pouring and early foam formation. The protein exhibits exceptional thermal stability during malting, mashing, and wort boiling (Evans and Hejgaard 1999, Jégou et al. 2001); however, partial unfolding induced by heat exposure promotes the exposure of hydrophobic residues, thereby enhancing its surface activity and membrane-forming capacity (Perrocheau et al. 2006; Mills et al. 2009). Consistent with this behaviour, LTP1 acts primarily during the initial stage of foam formation, facilitating bubble generation and stabilising newly formed foam films through rapid adsorption at the gas–liquid interface.

Proteomic analysis in the present study revealed an abundance of LTP1 in barley-based beers, whereas LTP1 was not detected, or was present only at trace levels, in all GF samples analysed. Although GF grains may contain other non-specific lipid transfer proteins (nsLTPs), their presence does not necessarily imply functional equivalence with barley LTP1. Like many plant nsLTPs, barley LTP1 adopts a compact, disulphide-stabilised helical fold; however, its brewing relevance has been specifically characterised. Barley LTP1 has a small hydrophobic cavity that can bind different lipid species, including fatty acids and acyl-CoA, and during malting and brewing it undergoes glycation, lipid adduction and heat-induced unfolding, changes that increase its amphiphilic character and promote adsorption at the air–water interface (Steiner et al. 2011). Therefore, the absence of barley LTP1 may represent the loss of a well-characterised brewing-responsive, interfacial-active protein in GF beers, although the foam-related functions of non-barley nsLTPs require further structural and interfacial characterisation.

### Mechanical reinforcement of the foam film by Protein Z

In contrast to the predominantly interfacial role of LTP1, Protein Z contributes mainly to the mechanical stabilisation and persistence of beer foam. As members of the serpin protein family, Protein Z isoforms are characterised by a β-sheet-rich fold capable of forming intermolecular interaction networks (Evans and Hejgaard 1999). Such structural features are well suited to reinforcing foam films by increasing elasticity and resistance to deformation, thereby slowing liquid drainage and bubble coalescence during foam retention (Evans and Hejgaard 1999; Niu et al. 2018).

In this study, two isoforms of Protein Z were consistently identified in barley-based beers, whereas no soluble serpin proteins were detected in any of the GF samples. This clear presence/absence pattern closely mirrored the experimentally observed differences in foam stability. Compared with storage or metabolic proteins commonly detected in GF beers, Protein Z possesses structural attributes that favour long-term interfacial reinforcement (Maeda et al. 2018). Its absence in GF systems likely compromises the mechanical strength of the foam film, leading to rapid collapse following initial formation.

### A dual-mechanism model for cooperative foam stabilisation

Based on the findings of the present study and previous literature on beer foam-active proteins, the results can be integrated into a conceptual model describing the complementary roles of LTP1 and Protein Z in barley beer foam formation and stabilisation (Figure 6). In this framework, LTP1 functions predominantly during the early stages of foam formation, where its strong interfacial activity promotes bubble generation and establishes the primary foam framework. Protein Z subsequently accumulates at the interface during the stabilisation phase, where it enhances film elasticity and mechanical resilience, thereby maintaining foam integrity over time. Together, these proteins act in a temporally and functionally complementary manner to produce a fine, stable, and persistent foam.

**Figure 6.**
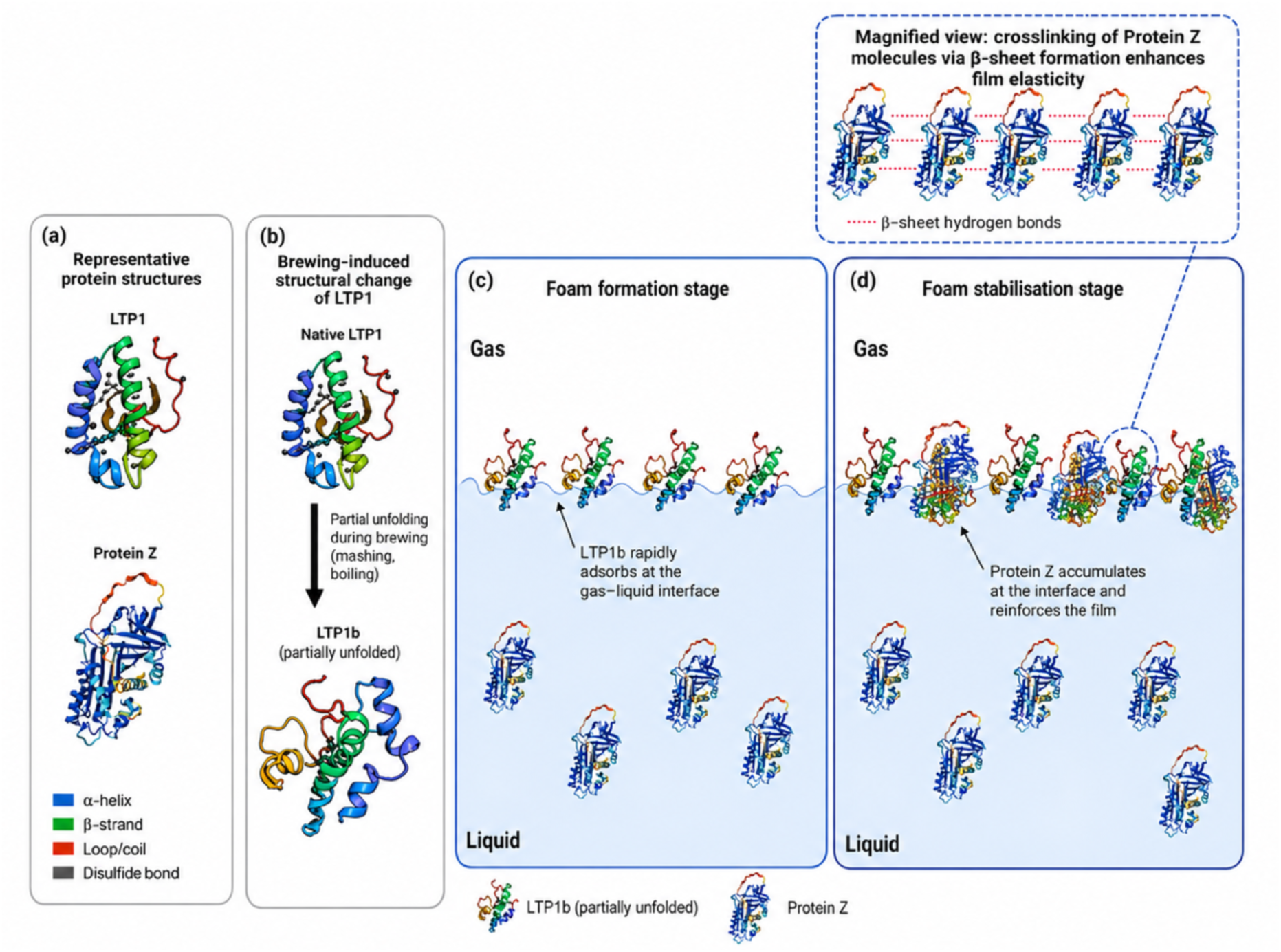

In GF beers, the protein network described above appears not to be able to form. The absence or trace amounts of LTP1 may reduce the availability of a well-characterised interfacial-active protein during the early stages of foam development, while the lack of detectable Protein Z may further limit the formation of a resilient stabilising protein film. Together with the lower overall soluble protein levels observed in GF samples, this loss of specific foam-associated proteins may help explain the reduced foam formation and rapid foam collapse observed in these beers. These findings suggest that beer foam quality is influenced not only by total soluble protein abundance, but also by the presence of proteins with suitable interfacial functionality.

### Limitations and future directions

While the present study provides new insights into the molecular determinants of beer foam stability, several limitations need to be acknowledged. First, the proteomic dataset was derived from a limited number of commercial beer samples for which complete information on recipe formulation, processing, and some other characteristics was not obtained. The observed differences in protein abundance and foam stability may therefore reflect differences beyond their gluten status, such as carbonation level, initial/final gravity, use of adjuncts, additives and processing aids, and mashing and fermentation conditions. Nevertheless, the comparison remains informative because the sampled beers represent commercially available barley-based and GF products, and the magnitude and consistency of the differences observed suggest that GF beers on the current market may face broader protein-related constraints on foam stability.

Second, protein identification was based on LC–MS/MS analysis of soluble fractions. Although this approach captured most surface-active proteins, it may have overlooked membrane-associated, highly hydrophobic, or low-abundance proteins that also contribute to interfacial behaviour.

Notably, among the three gluten-free samples (rice, sorghum, and the mixed-grain sample composed of millet, buckwheat, and rice), the mixed-grain sample yielded a markedly lower number of identifiable proteins. This may reflect analytical factors, such as extraction inefficiency, poor solubility of some grain proteins, or matrix-induced signal suppression, rather than a true absence of proteins. More broadly, differences in raw material form and recipe design across commercial beers, including malting status or the use of syrups and other adjuncts, may also influence the soluble protein profiles observed in this study.

Third, the proposed dual mechanism model proposed here to describe the cooperative function of LTP1 and Protein Z isoforms was inferred primarily from structural predictions and abundance correlations. Direct experimental evidence of physical interaction between these proteins remains limited. Future studies employing immunoprecipitation, surface plasmon resonance, or cross-linking mass spectrometry would be valuable to determine whether LTP1 and Protein Z isoforms form interfacial complexes during foam stabilisation. Complementary approaches such as interfacial rheology and microscopy could further elucidate the dynamic behaviour of these proteins at the air–liquid interface. In addition, supplementing GF beer systems with purified LTP1 or Protein Z could help assess their individual and combined contributions to foam formation and retention.

Finally, this study focused mainly on barley and three gluten-free beers derived from rice, sorghum, and a mixed-grain formulation. Expanding this framework to include a broader range of commercially relevant GF brewing grains and formulations, including millet-, buckwheat-, maize-, rice-, and sorghum-based beers, would provide a more comprehensive understanding of how grain-specific proteomes influence foam properties. Integrating proteomic with metabolomic data could also help identify compounds, such as Maillard reaction products, that may modulate the activity of foam-active proteins.

In summary, although several methodological and interpretative constraints exist, this study establishes a foundation for understanding the protein-based mechanisms underlying beer foam stability across diverse grain systems. Future research combining targeted proteomics, functional assays, and interfacial modelling will be essential to validate and refine the dual mechanism model proposed here.

## Conclusion

This study demonstrates that the reduced foam stability observed in GF beers is accompanied by lower soluble protein levels and substantial differences in protein composition. Barley-based beers contained higher protein concentrations and had higher abundances of LTP1 and serpin family members (Protein Z4 and Z7), which contribute to foam formation and retention, whereas GF beers showed markedly lower protein abundance and limited detection of these key protein groups.

The findings indicate that impaired foam performance in GF systems likely reflects a combination of low overall protein content and the absence of specific proteins that support interfacial adsorption and film stability. Consistent with previous literature, these patterns support a working model in which LTP1 is associated with early foam formation, whereas Protein Z is associated with later-stage foam stabilisation.

Together, these results provide a molecular framework for understanding the differences in foam persistence between barley-based and GF beers and offer a basis for improving foam stability through targeted formulation strategies.

## Author contributions (CRediT)

Zesun Lyu: Conceptualisation, Methodology, Investigation, Formal analysis, Data curation, Visualisation, Writing – original draft, Writing – review & editing.

Jacob Humphries: Conceptualisation, Methodology, Writing – review & editing.

Farhad Masoomi-Aladizgeh: Methodology, Validation, Supervision, Writing – review & editing.

Thomas H. Roberts**\***: Conceptualisation, Methodology, Supervision, Resources, Project administration, Funding acquisition, Writing – review & editing.

## Acknowledgements

The authors acknowledge the support of the School of Life and Environmental Sciences, University of Sydney, for access to laboratory facilities and equipment. The authors also thank the laboratory technical staff and Sydney Mass Spectrometry for their assistance during this study. This research received no specific grant from any funding agency in the public, commercial, or not-for-profit sectors.

## Conflict of interest

The authors declare that there are no conflicts of interest.

## Notes

### Competing Interest Statement

The authors have declared no competing interest.

