## Supplementary Tables S1-S5 for "Poor foam stability in gluten-free beers is associated with distinct protein composition compared with barley-malt beers"

**Table S1.** 10 most abundant proteins identified in the rice beer based on LFQ intensity.

**Rice Beer**

| Leading protein accession | Protein name | Intensity | Log2 Intensity |
|---|---|---|---|
| I1PWZ3 | Bifunctional inhibitor/plant lipid transfer protein/seed storage helical domain-containing protein | 65075 | 15.99 |
| T1T4Y4 | Glutelin | 41743 | 15.35 |
| Q6ERU3 | Glutelin type-B 5 | 17238 | 14.07 |
| A1YQH6 | Glutelin | 13387 | 13.71 |
| A0A0E0LII4 | Bifunctional inhibitor/plant lipid transfer protein/seed storage helical domain-containing protein | 11092 | 13.44 |
| W8NXS2 | Glyceraldehyde-3-phosphate dehydrogenase | 8787 | 13.10 |
| Q852L2 | Cupincin | 8184.9 | 13.00 |
| A0A063BWF5 | Phosphoglycerate kinase | 7215 | 12.82 |
| A0A6J2XGA7 | Enolase | 7069.1 | 12.79 |
| A0A8J5VEQ4 | Bifunctional inhibitor/plant lipid transfer protein/seed storage helical domain-containing protein | 6582.5 | 12.68 |

**Table S2.** 10 most abundant proteins identified in the sorghum beer based on LFQ intensity.

**Sorghum Beer**

| Leading protein accession | Protein name | Intensity | Log2 Intensity |
|---|---|---|---|
| A0A921U8M5 | Non-specific lipid-transfer protein | 295000 | 18.17 |
| A0A066XPY6 | PXA domain-containing protein | 13758 | 13.75 |
| A0A921UUK1 | HP domain-containing protein | 8723.7 | 13.09 |
| C5YQD9 | Calmodulin-binding domain-containing protein | 3305 | 11.69 |
| A0A066X9Q2 | tRNA (guanine(9)-N1)-methyltransferase | 927.99 | 9.86 |
| A0A066XNR0 | Kinetochore protein fta7 | 786.59 | 9.62 |
| A0A921U243 | Protein RDM1 | 771.41 | 9.59 |
| A0A921U5Q1 | Protein kinase domain-containing protein | 651.51 | 9.35 |
| A0A921S787 | Bifunctional inhibitor/plant lipid transfer protein/seed storage helical domain-containing protein | 531.04 | 9.05 |
| A0A194YN60 | RIN4 pathogenic type III effector avirulence factor Avr cleavage site domain-containing protein | 521.38 | 9.03 |

**Table S3.** 10 most abundant proteins identified in the mixed-grain beer based on LFQ intensity. The matched grain source is indicated for each protein identification.

**Mixed Grain Beer**

| Leading protein accession | Protein name | Matched grain source | Intensity | Log2 Intensity |
|---|---|---|---|---|
| K3YK93 | Non-specific lipid-transfer protein | Millet | 178770 | 17.45 |
| A0A3L6SLK5 | Non-specific lipid-transfer protein | Millet | 15276 | 13.90 |
| A0A3L6SB78 | Bifunctional inhibitor/plant lipid transfer protein/seed storage helical domain-containing protein | Millet | 12778 | 13.64 |
| A0A3L6Q3A5 | Non-specific serine/threonine protein kinase | Millet | 6319.5 | 12.63 |
| A0A3L6RH96 | 60S ribosomal protein L5-1 | Millet | 5597.8 | 12.45 |
| A0A3L6S733 | Non-specific lipid-transfer protein 2P-like | Millet | 4335.2 | 12.08 |
| D2CUB4 | Photosystem I P700 chlorophyll a apoprotein A1 | Buckwheat | 4235.3 | 12.05 |
| K3Z5B5 | Malectin-like domain-containing protein | Millet | 3564.3 | 11.80 |
| P83004 | 13S globulin basic chain | Buckwheat | 847.79 | 9.73 |
| H1ACZ4 | Early flowering 3 | Buckwheat | 837.23 | 9.71 |

**Table S4.** 10 most abundant proteins identified in the lager sample based on LFQ intensity.

**Barley (Lager)**

| Leading protein accession | Protein name | Intensity | Log2 Intensity |
|---|---|---|---|
| P07597 | Non-specific lipid-transfer protein 1 | 1116500 | 20.09 |
| Q40045 | D-hordein | 75557 | 16.21 |
| P06293 | Serpin-Z4 | 68798 | 16.07 |
| P11643 | Alpha-amylase/trypsin inhibitor CMd | 39840 | 15.28 |
| P28041 | Alpha-amylase/trypsin inhibitor Cma | 32824 | 15.00 |
| N1JH96 | Helix-loop-helix/DNA-binding domain protein | 30522 | 14.90 |
| A0A8I6W3I6 | Helix-loop-helix/DNA-binding domain protein | 26555 | 14.70 |
| Q546U1 | Barley dimeric alpha-amylase inhibitor (Bdai-1) | 22387 | 14.45 |
| A0A1E1LVR3 | Mannan endo-1,6-alpha-mannosidase | 21854 | 14.42 |
| A0A8I6WXV5 | BSD domain-containing protein | 21854 | 14.42 |

**Table S5.** 10 most abundant proteins identified in the pale ale sample based on LFQ intensity.

**Barley (Pale Ale)**

| Leading protein accession | Protein name | Intensity | Log2 Intensity |
|---|---|---|---|
| P07597 | Non-specific lipid-transfer protein 1 | 1502400 | 20.52 |
| P06293 | Serpin-Z4 | 101770 | 16.63 |
| A0A8I6XS48 | Bifunctional inhibitor/plant lipid transfer protein/seed storage helical domain-containing protein | 49882 | 15.61 |
| P28041 | Alpha-amylase/trypsin inhibitor Cma | 44517 | 15.44 |
| Q43492 | Serpin-Z7 | 29871 | 14.87 |
| P11643 | Alpha-amylase/trypsin inhibitor CMd | 26619 | 14.70 |
| Q9SES6 | Non-specific lipid-transfer protein | 26588 | 14.70 |
| A0A8I6W3I6 | Bifunctional inhibitor/plant lipid transfer protein/seed storage helical domain-containing protein | 22591 | 14.46 |
| A0A8I6XQ39 | Bifunctional inhibitor/plant lipid transfer protein/seed storage helical domain-containing protein | 19396 | 14.24 |
| Q5UNP2 | Non-specific lipid-transfer protein | 19013 | 14.21 |
